# Dynamics of Single-Base Editing: Theoretical Analysis

**DOI:** 10.1101/2023.05.08.539865

**Authors:** Vardan Hoviki Vardanyan, Qian Wang, Anatoly B. Kolomeisky

## Abstract

Recent experimental advances led to the development of DNA base editors (BEs) with a single-nucleotide precision that is critical for future progress in various scientific and technological fields. The molecular mechanisms of single-base discrimination, however, remain not well understood. Using a recently developed stochastic approach, we theoretically investigated the dynamics of single-base editing. More specifically, transient and mean times to edit “TC” motifs by cytosine BEs are explicitly evaluated for correct (target) and incorrect (bystander) locations on DNA. In addition, the effect of mutations on the dynamics of the single-base edition is also analyzed. It is found that for most ranges of parameters, it is possible to temporarily separate target and bystander products of base editing, supporting the idea of dynamic selectivity as a method of improving the precision of single-base editing. We conclude that to improve the efficiency of single-base editing, selecting the probability or selecting the time requires different strategies. Physical-chemical arguments to explain the observed dynamic properties are presented. The theoretical analysis clarifies some important aspects of molecular mechanisms of selective base editing.

## I. INTRODUCTION

One of the most striking recent developments that strongly affected multiple scientific and technological areas, ranging from chemistry and biology to engineering and medicine, is the discovery of experimental methods that allow for precise genome editing^1–3^. Originally based on utilizing clustered regularly interspaced short palindromic repeat (CRISPR) techniques, they have been significantly improved in recent years, revolutionizing multiple research fields.^4–9^ While a wide range of gene editing tools that utilized the CRISPR-Cas9 methods has been proposed,^4,10^ most of them require double-strand DNA breaks, frequently leading to unpredictable editing outcomes. Much more precise editing is achieved by base editors (BEs) that are constructed by fusing protein domains with specific enzymatic properties (for example, deaminases) and nickase Cas9 proteins that allow quick location to the proper site on DNA.^7,11–13^ In this case, the process is taking place without double-strand breaks in DNA, and this provides much more efficient and controllable genome editing.^6,9,11,12^

A wide spectrum of single-base editors has been reported in recent years.^12–16^ These biologically engineered systems allow for better editing precision and higher purity of products^17,18^. At the same time, the main problem of BEs remains the discrimination of identical bases in the activity windows (typically, 4-10 nucleotides) of these enzymatic complexes. In other words, it is challenging for the editor to modify only the specific base at the given site if identical bases can be found close to the target in the region labeled as the activity window. As a result, the bystander or both target and bystander nucleotides might be modified, negatively impacting the efficiency and precision of genome editing. While some improvements have been made by utilizing beneficial mutations in enzymatic domains, the discrimination issue has not been fully resolved yet.^14,19,20^ The main problem here is that the molecular mechanisms of the underlying processes remain not well clarified.

To assist in understanding the microscopic features of base editing and to rationally design more efficient BE systems, a theoretical approach that combines a discrete-state stochastic model with all-atom molecular dynamics has been proposed recently by one of us.^21^ It presented a minimal chemical-kinetic model that accounts for the most relevant chemical states of base editing and includes the possibility of target and bystander editing. More specifically, cytosine BE that converts cytosine (C) to thymine (T) has been considered.^14^ This process is shown schematically in Fig. 1. Some transition rates for the stochastic model have been estimated using various experimental observations,^14,20,22^ while the rest of parameters have been obtained using all-atom molecular dynamics simulations.^21^ The method of first-passage probabilities^23–25^ has been utilized then to characterize the features of base editing. This theoretical approach clarified some important molecular aspects of the process, and it proposed a set of general rules for designing BEs with improved editing selectivity. Importantly, the theoretical predictions of decreased bystander effect for specific mutations have been successfully verified by experiments.^21^

**FIG. 1:**
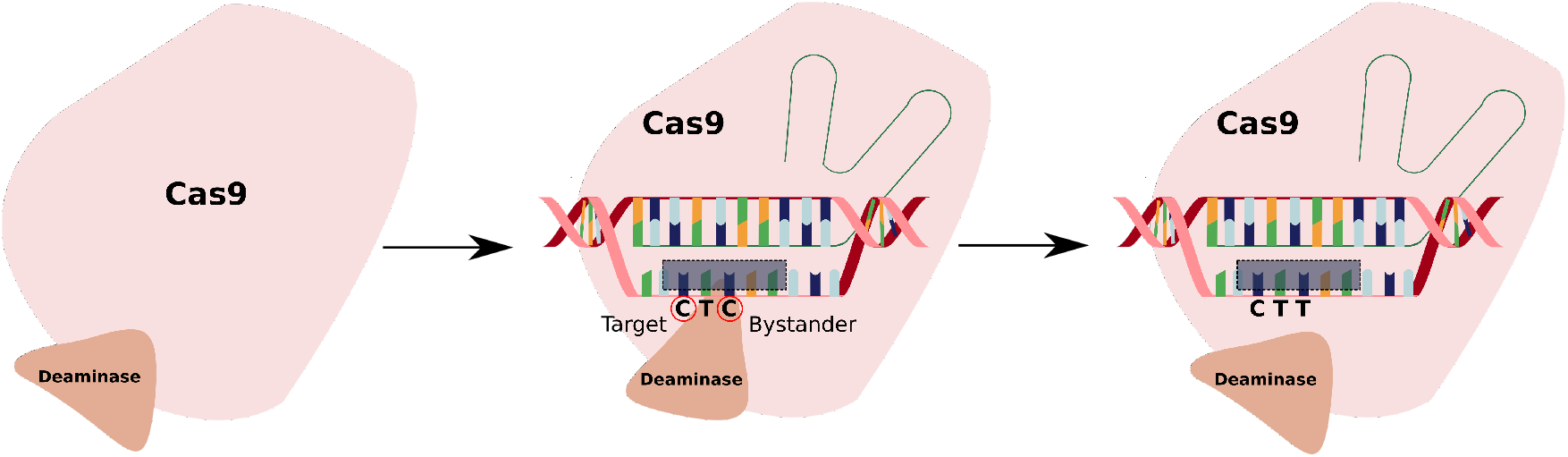
A schematic view of the functioning of the cytosine single-base editor. The grey area indicates the activity window of this BE.

Although this theoretical approach^21^ has provided some crucial information on the mechanisms of base editing, it has some limitations. This method mostly concerns with thermodynamic aspects of base editing since only the probabilities of different editing outcomes have been considered. The dynamics of base editing have not been effectively evaluated at all. However, different products might appear in the system at different times, and in real experiments, the most optimal conditions from the point of view of probability might not be realized because it might take too long to achieve them. Different times for different products also suggest that the selectivity of BEs might be improved by temporarily separating the products. The idea of improving the precision by collecting the products at different times is similar to the concept of dynamic selectivity.^26^ These observations raise several important questions. What is the dynamics of transient processes during the base editing? At what times do different products (target and bystander) appear in the system? Is it possible to separate them to enhance selectivity? And if yes, is the range of parameters for which the optimal dynamic selectivity might be observed correlated with the range of parameters when the bystander effect is reduced?

To answer these questions, we extend the original theoretical approach^21^ to explicitly evaluate the dynamics of transient processes, and calculate the mean editing times for different outcomes of single-base discrimination. This allows us to understand better the complex dynamics of single-base editing. Our analysis suggests that there are parameters at which the different products of base editing can be temporarily separated, exhibiting dynamic selectivity. Interestingly, it is found that the best conditions to dynamically separate the target editing do not correlate with the situations at which the probability of bystander editing is small. Physical-chemical arguments to explain these observations are presented. Our theoretical method allows for a better understanding of microscopic processes associated with single-base editing in DNA.

## II. THEORETICAL METHOD

To investigate the dynamics, we utilize a minimal chemical-kinetic model already developed to describe the process of genome editing of the EGFP site 1 by A3A-BE3 cytosine BE.^21^ It is schematically presented in Fig. 2. Note that although the total number of states in this chemical-kinetic description is relatively large (total 15 states), they describe four major pathways in this system that cannot be neglected if one aims to correctly analyze genome editing by BEs.^21^

**FIG. 2:**
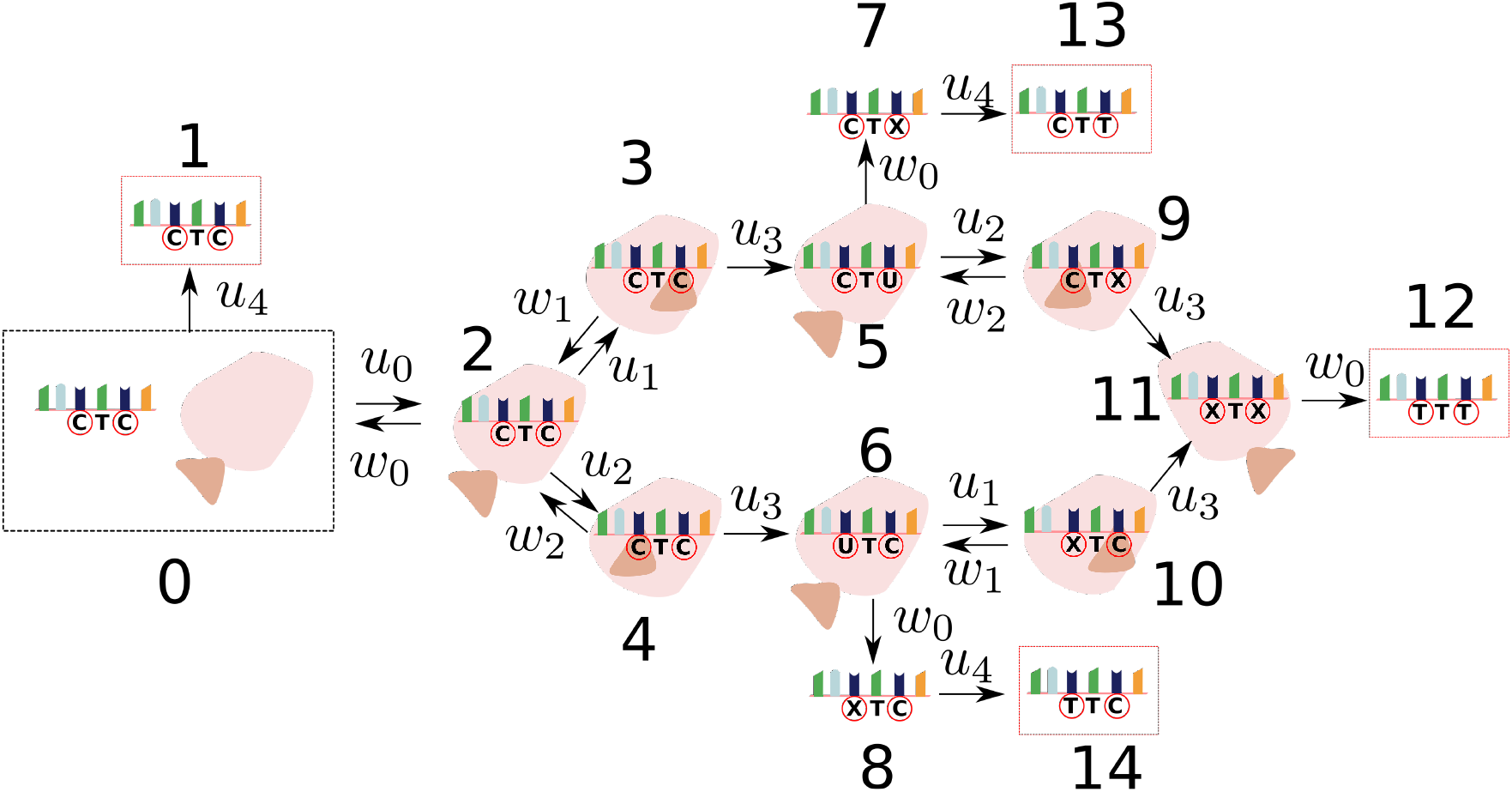
Chemical-kinetic model to describe single-base transformations by cytosine BE. The details of the process are discussed in the text. The system starts in the state 0, and there are four possible outcomes. The state 1 corresponds to no editing events, state 12 describes the editing at both target and bystander sites, state 13 corresponds to the target-site editing (the desired outcome), and state 14 describes the bystander editing. The nucleotides labeled “C” or “T” correspond to cytosine or thymine, respectively, while the nucleotide labeled “X” corresponds to uridine or thymine.

The base-editing process starts in the state 0, and then there are two possible transitions out of this state. The BE can change into inactive conformation (transition 0 → 1) with a rate *u*_4_, which corresponds to the situation when no editing events happened. But if the BE stays active, its Cas9 domain can associate with the single-stranded DNA segment with a rate *u*_0_ (transition 0 → 2). This process is reversible (2 → 0), and the corresponding dissociation rate is *w*_0_. Then there are two possibilities. The deaminase domain of the BE can bind to the target nucleotide with a rate *u*_1_ (2 → 3), or it can bind to the bystander cytosine with a rate *u*_2_ (2 → 4). Since the target and bystander nucleotides are chemically identical and spatially very close, it is reasonable to assume that *u*_1_ = *u*_2_.^21^ Both of these processes are reversible and the backward reaction rates are *w*_1_ (3 → 2) and *w*_2_ (4 → 2). In the next step, the enzymatic reaction converts cytidine (C) into uridine (U) with the same rate *u*_3_ for the target site (3 → 5) and the bystander site (4 → 6). After that, from the states 5 or 6 there are two possible chemical pathways. If the Cas9 domain dissociates from DNA, the uridine will be modified into thymidine by a DNA repair mechanism (transitions 5 → 7 → 13 or 6 → 8 → 14). State 13 corresponds to the product of correct editing (target CTT), while state 14 corresponds to the wrong product (bystander TTC). However, if Cas9 domain stays longer on DNA it can also modify the neighboring nucleotides (transitions 5 → 9 → 11 → 12 or 6 → 10 → 11 → 12) with the corresponding transition rates: see Fig. 2. These events result in the wrong product where both target and bystander nucleotides are edited (TTT, state 12).

The transition from the state 11 to the state 12 should also have two steps with the rates *w*_0_ and *u*_4_, similarly to 5 → 7 → 13 and 6 → 8 → 14. But because for the experimentally estimated rates it was found that *w*_0_ ≪ *u*_4_,^21^ this transition can be viewed as a single step with the limiting rate *w*_0_. This approximation does not affect any of the results of our calculations.

One should also note that cytosine and thymine nucleotides are labeled as “C” and “T”, respectively (Fig. 2). At the same time, the nucleotide labeled as “X” represents both uridine and thymine, reflecting the fact that it is not known when the chemical transformation of uridine into thymine is taking place.

To obtain dynamic properties of single-base editing, we utilize a method of first-passage probabilities that has been widely explored for investigating stochastic processes in chemistry, physics, and biology.^21,23–25,27–30^ More specifically, we explicitly evaluate not only the probabilities of different editing outcomes, which was already accomplished before,^28^ but also mean editing times to obtain different products. To simplify our calculations, we notice that some of the transitions are irreversible (see Fig. 2), allowing us to consider the editing process (namely, reaching the states 12, 13, and 14) as a sequence of two events. In the first event, the system reaches the intermediate states 5 or 6 starting from the state 0, and in the second event, the final editing states are achieved after initiating in the state 5 or 6: see Fig. 2. This allows us to understand better transient processes in base editing.

Let us consider first processes that start in the state 0 and end in the state 5. One can introduce a function *F*_*j*_(*t*) defined as a probability density to reach the state 5 at time *t* for the first time if at *t* = 0 the system started in the state *j* (*j* = 0, 2, 3, 4). The temporal evolution of these fist-passage probability functions is given by a set of backward master equations,^21,23,24,29^

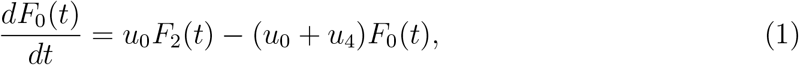

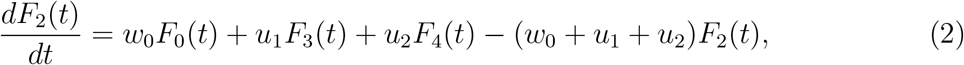

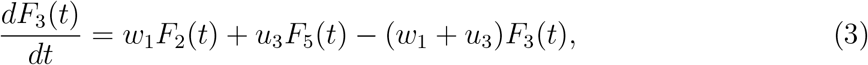

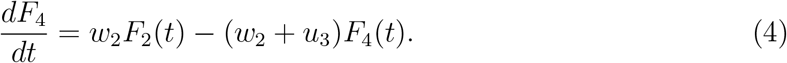

In addition, there is a boundary condition *F*_5_(*t*) = *δ*(*t*), which has a physical meaning that if the system starts in the state 5 is process is immediately accomplished. These equations also assume that if the system reaches the states 1 or 6 these are unsuccessful events, i.e. *F*_1_(*t*) = *F*_6_(*t*) = 1 at all times.

The system of backward master equations can be solved by utilizing Laplace transformation of the first-passage probability functions, 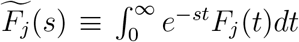, which allows us to modify the original differential equations into a system of algebraic equations,

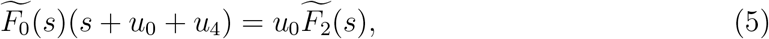

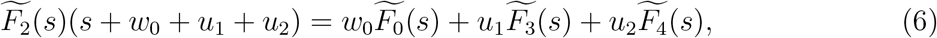

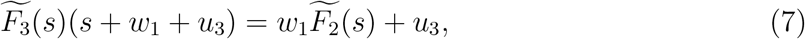

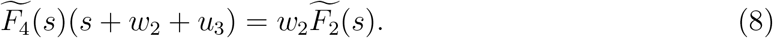

These equations can be exactly solved to obtain the explicit expressions for 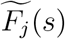, from which one can estimate the dynamic properties of the process. More specifically, we evaluate the probability of the event (*P*_0→5_) and the mean time before this happens (*T*_0→5_). It can be shown that

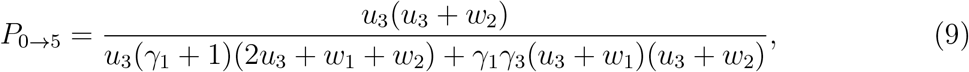

and

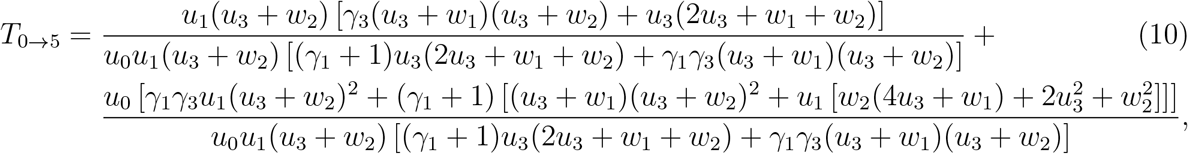

where parameters *γ*_1_, *γ*_2_ and *γ*_3_ are given by^21^

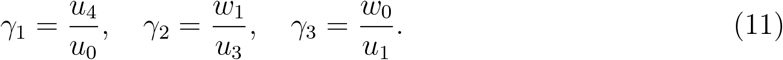

Similar analysis can be done for another process in the first step of base editing, 0 → 6 (see Fig. 2). One can derive that the probability of the event (*P*_0→6_) and the mean time before this happens (*T*_0→6_) are given by the following expressions,

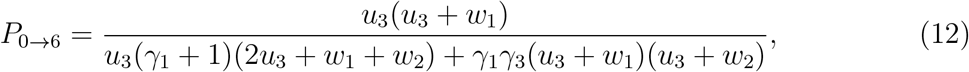

and

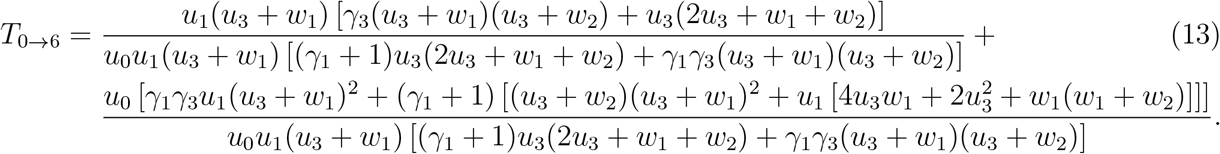

In Fig. 3a, we present the mean times for the first stage of base editing as a function of the binding rate of the deaminase domain to the nucleotide. As one can see, increasing the binding rate *u*_1_ lowers the times to reach these states, which is expected. However, what is less expected is that the events along the less probable pathway are faster (note that *P*_0→6_ ≪ *P*_0→5_ - see caption to Fig. 3). This can be explained using the following arguments. Because *w*_2_ *> w*_1_ (see Fig. 2), the system is rarely able to reach the state 6 from the state 4. Only fast transitions are able to do it. As a result, we have for the mean times *T*_0→6_ < *T*_0→5_, although the probabilities of these events are quite small, *P*_0→6_ ≪ *P*_0→5_.

**FIG. 3:**
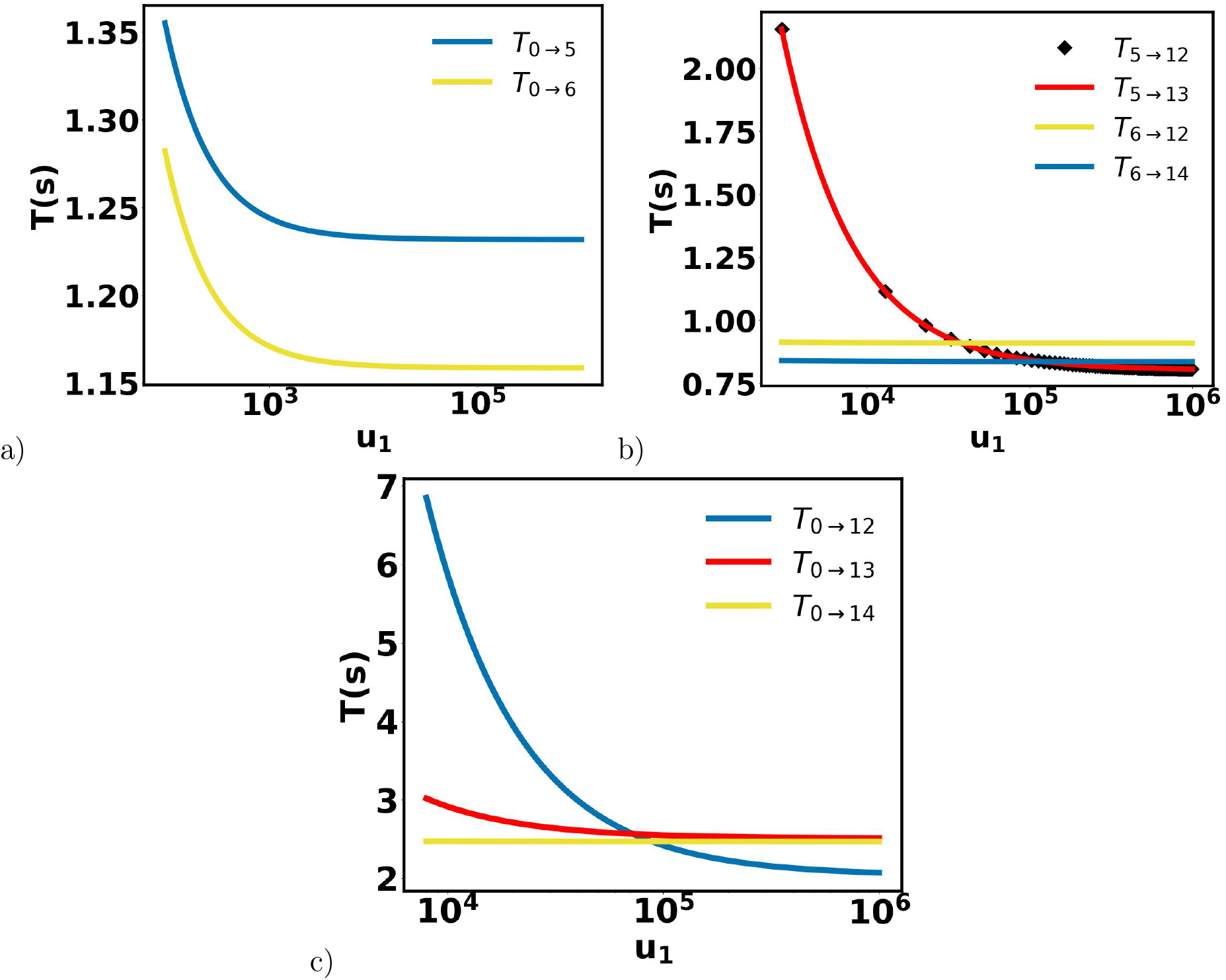
Dynamics of different base-editing processes as a function of the binding rate of deaminase domain to the nucleotide *u*_1_. The following parameters have been used in calculations:^21^ *u*_0_ = 1 s^−1^, *u*_3_ = 1.1 s^−1^, *u*_4_ = 2.1 s^−1^, *w*_0_ = 2.9 * 10^−5^ s^−1^, *w*_1_ = 12.54 s^−1^ and *w*_2_ = 5059 s^−1^. a) Mean times for 0 → 5 and 0 → 6 processes in the first stage of base editing. The corresponding probabilities are *P*_0→5_ = 0.321635 and *P*_0→6_ = 0.000867. b) Mean times for 5 → 12, 5 → 13, 6 → 12 and 6 → 14 processes in the second stage of base editing. The corresponding probabilities are *P*_5→12_ = 0.882299, *P*_5→13_ = 0.117701, *P*_6→12_ = 0.999641, and *P*_6→14_ = 0.000359. c) Mean times for the overall target editing 0 → 13, bystander editing 0 → 14 and double editing 0 → 12. The corresponding probabilities are *P*_0→13_ = 0.0378568, *P*_0→14_ = 3.11661 * 10^−7^ and *P*_0→12_ = 0.284645.

The second stage of the base editing process starts in the states of 5 or 6 (Fig. 2), and there are three final outcomes. Editing the target nucleotide leads to the state of 13, editing the bystander nucleotide produces the state of 14, and editing both target and bystander nucleotides yields the state of 12. The first-passage analysis for these processes can be done in a similar way done above for the processes in the first step of base editing. We then obtain the probabilities,

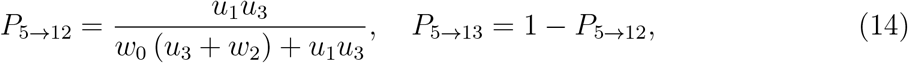

and

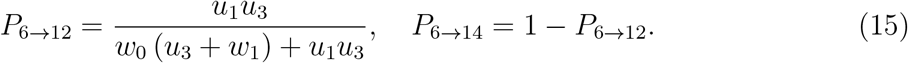

For the mean times for these events, the results are

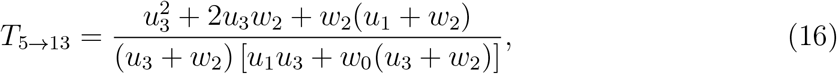

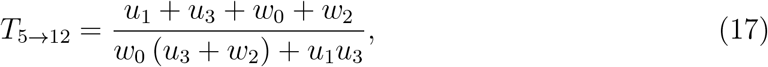

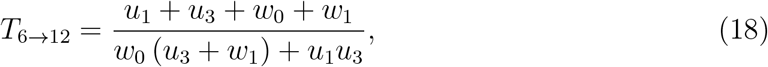

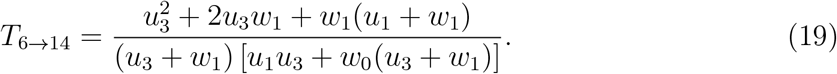

The results for dynamics in the second step of base editing are shown in Fig. 3b. Increasing the binding rate *u*_1_ accelerates the processes that start in the state 5, while there is a much smaller effect on the processes starting from the state 6. Again, the least probable process 0 → 14 is the fastest for most ranges of parameters, although for very fast binding rates the processes that start in the state 5 become slightly faster.

Now we can combine the analysis of the first and second steps of base editing to obtain the dynamic properties for the overall process. The probability and the mean time to edit the target nucleotide can be evaluated as

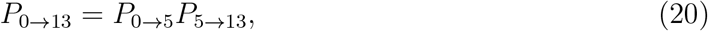

and

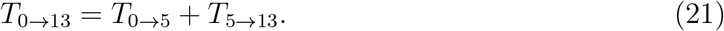

Similarly, one can derive for the editing of bystander nucleotide,

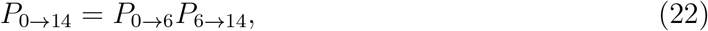

and

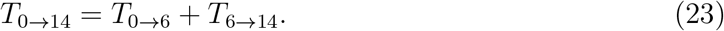

For the process of editing both target and bystander nucleotides, it can be shown that

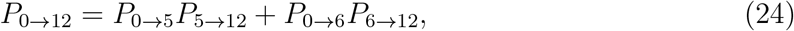

and

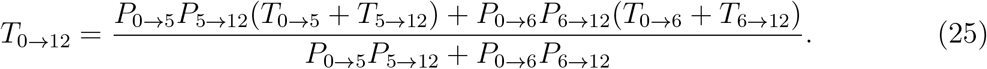

The physical meaning of this expression is the following. There are two pathways to reach the state 12 from the state 0. The factor 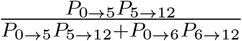 gives the probability for the system to choose pathway 0 →5 →12, and the factor 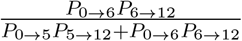 is the probability to follow the pathway 0 → 6 → 12. The explicit expressions for all dynamic properties for the overall base editing can be easily obtained from Eqs. (12)-(19).

Fig. 3c illustrates the overall dynamics of base editing for different processes. In all cases, increasing the rate *u*_1_ accelerates the dynamics, although the amplitude is not the same for different pathways. The strongest effect is observed for the processes of editing both bystander and target nucleotides (0 → 12). The weaker effect is observed for editing the target nucleotide (0 → 13), and there is almost no effect for editing the bystander nucleotide (0 → 14). But the important result is that the mean times to obtain different products of base editing in most ranges of parameters are not the same, suggesting that the products can be temporarily separated. This is the main reason to explore the idea of dynamic selectivity for improving the precision of single-base editing.

To optimize the performance of single base editors, the most common approach is to explore various mutations in the deaminase domain.^14,19–21^ Then the effect of mutation can be viewed as a perturbation in the energy of the base-editing process. In the simplest approximation, which however was also supported by some recent experimental data,^21^ it was argued that the effect will mostly occur in the dissociation steps of the deaminase from the DNA chain, leading to the following changes in unbinding transition rates,

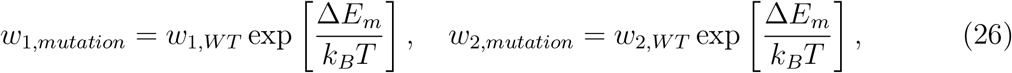

where Δ*E*_*m*_ is the free-energy difference in the deaminase unbinding process for the system with mutation in comparison with the wild-type (WT) case. When Δ*E*_*m*_ is positive the unbinding process is happening faster, while for Δ*E*_*m*_ < 0 it is more suppressed. Recent investigations have shown that for A3A-BE3 editing system the mutations that lead to the perturbations Δ*E*_*m*_ ≃ 5 − 7 *k*_*B*_*T* exhibit the decreased bystander effect, i.e., the probability of target editing in comparison with other outcomes is maximal at these conditions.^21^

The effect of mutations can be easily quantified in our theoretical approach. One can analyze the ratio of probabilities *P*_0→13_*/P*_0→14_, which specifies how more probable the target editing over the bystander editing, and the ratio of probabilities *P*_0→13_*/P*_0→12_, which measures how the target editing is more probable in comparison with editing both target and bystander nucleotides. The results for different free-energy perturbations associated with different mutations are presented in Fig. 4. One can see that the target editing (0 → 13) is much more probable that the bystander editing (0 → 14) for most ranges of parameters, but the largest effect is achieved for the systems with Δ*E*_*m*_ ≃ 0. Mutations associated with strongly negative or strongly positive free-energy changes decrease the advantage of the target editing. The situation is very different when we compare with the double editing events. For the mutations with Δ*E*_*m*_ *<* 0, it is more probable to obtain the products with editing in both target and bystander nucleotides. Only for strongly positive perturbations (Δ*E*_*m*_ ∼ 5−10 *k*_*B*_*T*), the target editing becomes more preferred. These are the conditions of reduced bystander effect, as one can also see from the ratio *P*_0→13_*/*[*P*_0→12_ + *P*_0→14_] presented in Fig. 4. This quantity describes the relative probability of obtaining the desired target editing in comparison with other undesired outcomes.

**FIG. 4:**
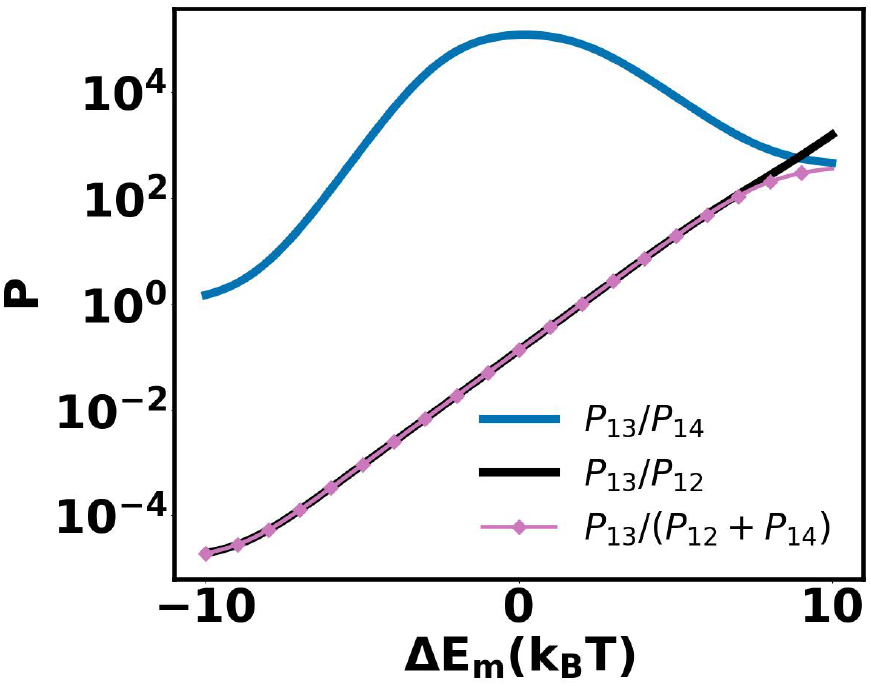
Ratio of probabilities of different outcomes of base editing as a function of free-energy perturbation due to mutations. The following parameters have been used in calculations:^21^ *u*_0_ = 1 s^−1^, *u*_3_ = 1.1 s^−1^, *u*_4_ = 2.1 s^−1^, *w*_0_ = 2.9 * 10^−5^ s^−1^, *w*_1_ = 12.54 s^−1^ and *w*_2_ = 5059 s^−1^.

To understand which editing products are coming first, in Fig. 5 we present the results of explicit calculations for the ratios of mean editing times for different binding rates *u*_1_ as a function of free-energy perturbations associated with possible mutations. More specifically, *T*_0→13_*/T*_0→14_, which describes how faster the target nucleotide is edited in comparison with the bystander nucleotide, and *T*_0→13_*/T*_0→12_, which describes how faster the target nucleotide is edited in comparison with editing both nucleotides, are calculated: see Fig. 5. For slow binding rates *u*_1_ (Fig. 5a), it is usually faster to edit the bystander nucleotide, although the probability for such events is quite low (compare with Fig. 4). As we already argued, this is because only the fastest events will go along the pathway 0 → 14, but they are quite rare. Increasing the binding rate *u*_1_ (Figs. 5b and 5c) changes the situation. It is still faster to edit the bystander nucleotide for Δ*E*_*m*_ < 0, while the desired target editing is faster for mutations associated with Δ*E*_*m*_ > 0.

**FIG. 5:**
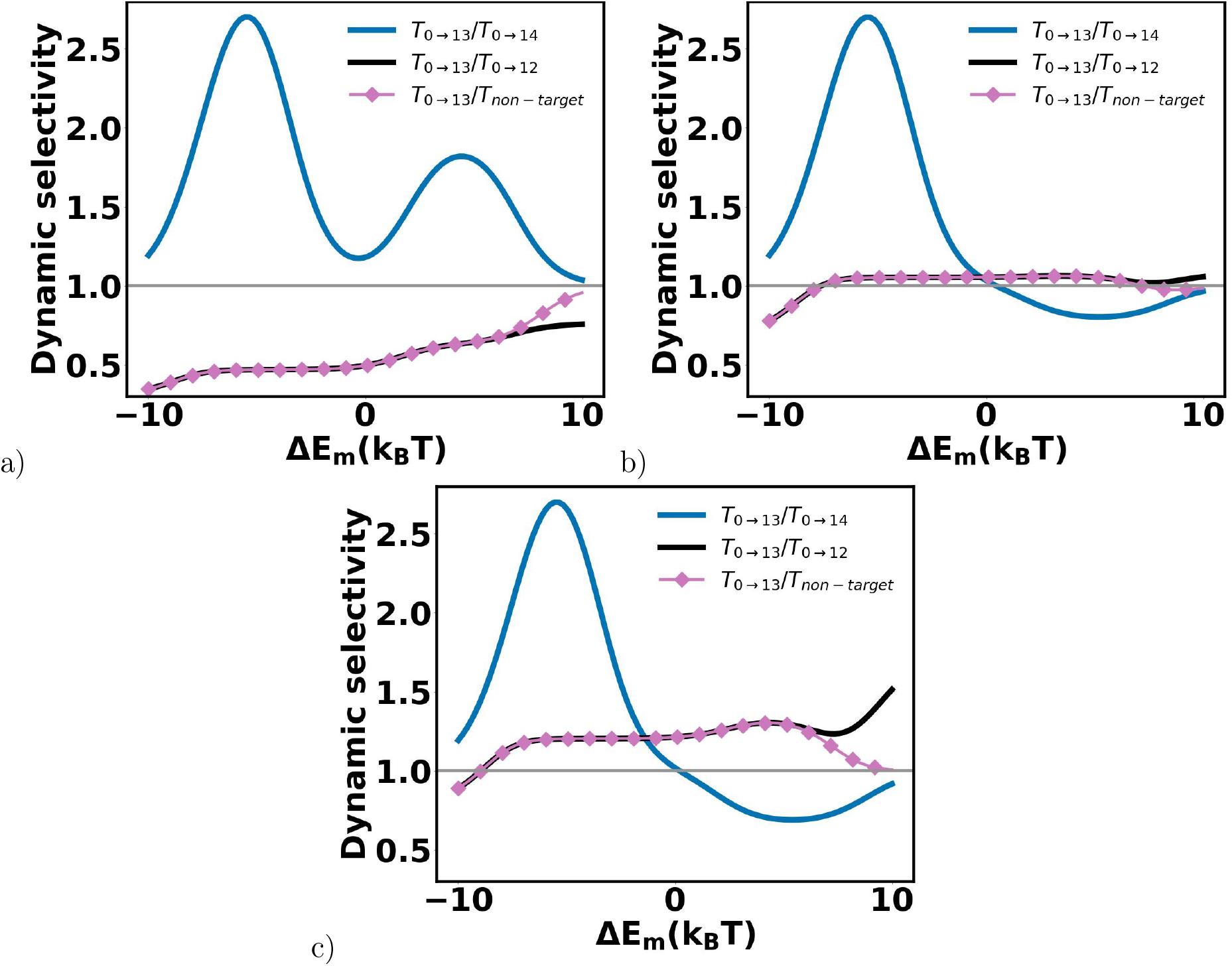
Ratios of mean editing times as a function of free-energy perturbation due to mutations. The following parameters have been used in calculations:^21^ *u*_0_ = 1 s^−1^, *u*_3_ = 1.1 s^−1^, *u*_4_ = 2.1 s^−1^, *w*_0_ = 2.9 * 10^−5^ s^−1^, *w*_1_ = 12.54 s^−1^ and *w*_2_ = 5059 s^−1^. a) For unbinding rate *u*_1_ = 10^4^ s^−1^. b) For unbinding rate *u*_1_ = 10^5^ s^−1^. c) For unbinding rate *u*_1_ = 10^6^ s^−1^.

Comparing the dynamics of target editing with editing in both locations, as quantified by the ratio *T*_0→13_*/T*_0→12_, one can see that for the slow binding rates *u*_1_ it is faster to edit the target nucleotide (Fig. 5a). But increasing the rate *u*_1_ makes both times scales comparable (Fig. 5b), and the target editing becomes even slightly slower for faster binding rates *u*_1_: see Fig. 5c.

Different editing times suggest that the precision of the single-base discrimination process can be improved by collecting the products at different times. We can quantify the conditions when such dynamic selectivity can lead to the most optimal performance. For this purpose, we define the mean time to obtain non-target undesired products,

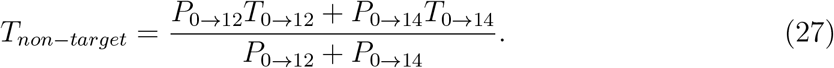

Then the ratio *T*_0→13_*/T*_*non*−*target*_ will provide a measure of dynamic selectivity in the single-base editing process. This quantity is plotted in Fig. 5. It is clear that the most efficient single-base discrimination occurs when *T*_0→13_*/T*_*non*−*target*_ deviates significantly from unity. One can see that the best selectivity can be achieved for slow binding rates *u*_1_ (Fig. 5a), and it is slightly better for the mutations with Δ*E*_*m*_ *<* 0. However, increasing the binding rate *u*_1_ effectively eliminates the possibility of temporal separation (Fig. 5b). But further increase in the rates *u*_1_, starts to improve the selectivity but not too much. One could also observe that the mean time to obtain any non-target products for most ranges of parameters is close the mean time to edit both target and bystander nucleotides (*T*_*non*−*target*_ ≈ *T*_0→12_). This is because the editing of only the bystander nucleotide is quite fast and it has a very low probability.

Another important result of our theoretical calculations is that the conditions of the highest probability and the best dynamic selectivity for the desired target single-base editing generally do not correlate. In other words, the temporal separation at the conditions of the lowered bystander effect (Δ*E*_*m*_ ≃ 5 − 10 *k*_*B*_*T*) cannot be efficiently utilized. This important observation underlies the difference between thermodynamic and kinetic controls of single-base editing processes.

## III. SUMMARY AND CONCLUSIONS

In this paper, we presented a theoretical analysis of processes that are taking place during single-base editing. A specific system of cytosine BE has been considered. Using a minimal chemical-kinetic model together with the first-passage probabilities method, we explicitly evaluated the dynamic properties, precision, and selectivity of base editing. Our theoretical method allowed us to evaluate the probabilities and mean times of different editing outcomes.

In addition, the role of mutations in optimizing the performance of single-base editors is quantitatively estimated for experimentally relevant ranges of parameters. It is found that increasing the association rate of the enzymatic domain accelerates all editing processes, but it also lowers the degree of dynamic selectivity. Our calculations also show that mutations that increase the dissociation rates of enzyme domains provide the best conditions for the reduced bystander effect at which the probability of target editing is maximal. However, the best temporal separation can be achieved for mutations that lower these dissociation rates. Furthermore, we presented a detailed description of transient processes during base editing, allowing us to understand better the molecular mechanisms of underlying processes. Such information can help to rationally develop the most efficient single-base editing systems for various applications. The most important conclusion from our theoretical analysis is that there are different requirements to optimize the functioning of single-base editors from probabilistic or dynamic aspects of the process.

Although our theoretical approach has been able to obtain a comprehensive dynamic picture of base editing processes, it is important to discuss its limitations. The utilized chemical kinetic model is rather very simplified, and many more biochemical transitions are neglected. In addition, some chemical transitions are assumed to be irreversible while in reality, this might not be the case. However, despite these limitations, the most important advantage of our theoretical method is that it can provide specific quantitative predictions that can be experimentally tested. It will be important to investigate more advanced models and do more experimental studies in order to clarify the microscopic picture of precise single-base discrimination.

## ACKNOWLEDGEMENT

ABK acknowledges the supported form the Welch Foundation (C-1559), from the NIH (R01 HL157714-02), from the NSF (CHE-1953453), and from the Center for Theoretical Biological Physics sponsored by the NSF (PHY-2019745). WQ thanks the funding support from the National Natural Science Foundation of China (32000882).

## Notes

### Competing Interest Statement

The authors have declared no competing interest.

